# Amplification-Free Detection of Antibiotic Resistance in *Enterococcus faecium* using PNA-FISH

**DOI:** 10.64898/2026.04.24.720744

**Authors:** Jae-Kyeong Im, Seongmin Yun, Brian Choi, Sungho Kim, Joo H. Kang, Taejoon Kwon, Hajin Kim

## Abstract

Vancomycin-resistant *Enterococcus faecium* (VREfm) is a major nosocomial pathogen, with antibiotic resistance mediated by the *vanA* and *vanB* operons. Rapid and accurate detection of antibiotic resistance is critical for the timely treatment of bacteremia and sepsis. Although imaging-based approaches using fluorescence *in situ* hybridization (FISH) provide a potential diagnostic solution, detecting mRNAs of antibiotic resistance genes (ARGs) in individual cells remains particularly challenging due to their low copy number and transient expression. Here, we present a peptide nucleic acid (PNA)-FISH method for direct detection of *vanA-* and *vanB*-associated resistance in individual VREfm cells. A universal probe targeting the conserved region across vancomycin resistance genes and a set of probes exclusively targeting the *vanB* gene were designed. The universal probe showed increased fluorescence in the *vanA*-genotype strain upon vancomycin or teicoplanin treatment, and in the *vanB*-genotype strain upon vancomycin treatment. In contrast, *vanB*-specific probes showed increased fluorescence exclusively from the *vanB*-genotype strain upon vancomycin treatment, confirming their specificity to the *vanB* gene. Efficient cellular penetration and strong hybridization of PNA probes enabled efficient and accurate detection of antibiotic-resistant bacterial cells, even under a wide-field fluorescence microscope. No detectable signals above background were observed in other major bacterial species associated with bacteremia and sepsis. These findings demonstrate robust detection of antibiotic-resistant cells in mixed microbial populations. When integrated with microbe-capturing techniques, this method may support culture-free detection of antibiotic resistance without nucleic acid amplification or sequencing, with the potential to reduce diagnostic turnaround time.

## 1. INTRODUCTION

*Enterococcus faecium* (*E. faecium*) has emerged as a major nosocomial pathogen due to its remarkable adaptability and frequent acquisition of multidrug resistance (1–4). Genomic studies have shown that hospital-adapted lineages of *E. faecium* have evolved through extensive acquisition of mobile genetic elements and antibiotic resistance determinants (5). These mobile genetic elements facilitate both the rapid accumulation of resistance within *E. faecium* and the dissemination of antimicrobial resistance across hospital-associated *E. faecium* lineages (6). Notably, vancomycin resistance determinants carried by VREfm can be horizontally transferred to other Gram-positive pathogens. Several clinical cases of vancomycin-resistant *Staphylococcus aureus* (VRSA) have been reported in which methicillin-resistant *S. aureus* (MRSA) acquired the *vanA* operon from VREfm during co-infection or co-colonization (7,8). Among available therapeutic options, glycopeptide antibiotics such as vancomycin have long been regarded as last-line agents (9). However, the widespread dissemination of VREfm has become a critical global health concern (2,10).

Resistance is predominantly mediated by horizontally acquired clusters of ARGs, known as the *vanA* and *vanB* operons, which encode enzymes that remodel peptidoglycan precursors from the canonical D-Ala-D-Ala to D-Ala-D-Lac, thereby abolishing vancomycin binding (11,12). Although *vanA*- and *vanB*-type resistance account for most clinical isolates, their antibiotic susceptibility profiles differ: *vanA* typically confers resistance to both vancomycin and teicoplanin, whereas *vanB* mediates variable resistance to vancomycin while retaining susceptibility to teicoplanin (13,14). Distinguishing these genotypes has a direct therapeutic impact.

Current diagnostic workflows for VREfm rely on culture-based antimicrobial susceptibility testing or molecular PCR assays (15–18). Culture-based methods are time-consuming, delaying clinical decision-making, and PCR requires nucleic acid extraction and may not directly reflect gene expression (19). PNA-FISH offers a promising alternative to these approaches. PNA probes exhibit superior cellular penetration, high affinity and specificity for complementary RNA sequences due to their high sensitivity to mismatches, and resistance to nucleases (20). Importantly, PNA-FISH enables direct analysis of clinical samples without culture, thereby reducing diagnostic turnaround time (21).

In this study, we designed a universal PNA probe targeting a conserved region shared by *vanA* and *vanB*, and a set of probes specifically targeting *vanB*. We evaluated their ability to distinguish between *vanA*- and *vanB*-genotype *E. faecium* under antibiotic induction. In addition, we investigated the feasibility of multiplex detection using probes labeled with spectrally distinct fluorophores. Our results demonstrate that the PNA-FISH method enables rapid and accurate detection and differentiation of vancomycin resistance genotypes at the single-cell level, supporting its potential utility for guiding antibiotic therapy and infection control.

## 2. MATERIALS AND METHODS

### 2.1. Bacterial culture and antibiotic treatment

All bacterial species (*Enterococcus faecium* [NCCP 11522, NCCP S16 (National Culture Collection for Pathogens)], vancomycin-susceptible *Enterococcus faecium* [KCTC 13225 (Korean Collection for Type Cultures)], *Escherichia coli* [NCCP 2571], Staphylococcus aureus [KCCM 12103 (Korean Culture Center of Microorganisms)], *Klebsiella pneumoniae* [NCCP 16285], *Pseudomonas aeruginosa* [NCCP 2641, PAO1]) were streaked onto fresh LB agar plates and incubated overnight at 37°C. A single colony of each strain was inoculated into 3 mL LB medium and grown at 37°C with shaking. Bacterial growth was measured every 30 min by measuring optical density at 600 nm (OD600) using a UV-Vis spectrometer (NanoDrop 2000c, Thermo Fisher Scientific). Antibiotic treatment was initiated when OD600 reached 0.1-0.2. Vancomycin and teicoplanin (Sigma-Aldrich) were dissolved in ultrapure water at 10 mg/mL and added directly to growing LB medium. Final concentrations were 32 μg/mL for vancomycin and 16 μg/mL for teicoplanin.

### 2.2. Preparation of poly-L-lysine-coated plate

Black 96-well plates (SPL Life Sciences, South Korea) were sonicated in 1 M nitric acid for 30 min at room temperature. Plates were washed three times with ultrapure water and dried with nitrogen gas. The plates were then incubated with 0.01% poly-L-lysine (PLL) solution for 30 min at 37°C, followed by washing and drying. Coated plates were wrapped with parafilm and stored at 4°C until use.

### 2.3. Design of PNA-FISH probes

The sequences of *vanA* and *vanB* from the comprehensive antibiotic resistance database (CARD) were used to design the PNA FISH probes (22). For van-Univ probe, 15-base candidate sequences were generated from the conserved regions between *vanA* and *vanB*. For vanB-1, 2, 3 probes, 15-base candidate sequences were generated from the *vanB*-specific regions. Probes that meet the criteria for purine content, GC content, length of purine stretch, and self-complementarity were selected: purine content ≤ 50 %, guanine content ≤ 35 %, purine stretch length < 6 bp and passing both reverse complementary 7-mer and 3-mer self-complementarity checks. Any probe sequence with gapless alignment to the genomic DNA of *E. faecium* with a 2-base mismatch allowance was excluded. Then the sequences that minimize cross-hybridization to non-target bacterial species were selected.

### 2.4. PNA-FISH procedure

A 7 µL aliquot of bacterial suspension was applied to the center of each well of the PLL-coated plate and incubated for 5 min at room temperature. Cells were fixed with 200 µL of 3.7% formaldehyde for 30 min, followed by three washes with nuclease-free water. Cells were permeabilized with 100 µL ice-cold methanol at –20°C for 10 min and washed again. Hybridization was performed in 75 µL solution containing 10% (w/v) dextran sulfate, 10 mM NaCl, 30% formamide, 0.1% (w/v) sodium pyrophosphate, 0.1% Triton X-100, 50 mM Tris-HCl (pH 7.5), 0.2% (w/v) polyvinylpyrrolidone, 0.2% (w/v) Ficoll, 5 mM EDTA, and 200 nM PNA probe (Panagene, South Korea) for 1 hr at 55°C in a humidified chamber. Following hybridization, samples were washed twice in buffer containing 20% formamide, 5 mM Tris-base (pH 10, Biosesang, South Korea), and 15 mM NaCl at 55°C for 15 min to remove unbound probes.

### 2.5. Image acquisition and statistical analysis

Confocal images were acquired using a Zeiss LSM 980 microscope (Carl Zeiss AG, Germany) equipped with a 20× objective (Carl Zeiss, NA = 1.0). Excitation wavelengths of 561 nm and 633 nm were used for the Cy3 and Cy5 channels, respectively. Cell boundaries were detected from the Cy5 channel images, and fluorescence intensities in the Cy3 channel were quantified within the segmented regions after background subtraction. Wide-field images were acquired by THUNDER Imager (Leica Microsystems, Germany), using a 20× objective (Leica Microsystems, NA = 0.8). 440 nm and 550 nm excitation sources were used for Hoechst and Cy3 channels, respectively. Instant computational clearing (ICC; Leica Microsystems) was applied to wide-field imaging to reduce background signals. ICC estimates a baseline of background mainly originating from out-of-focus signals and subtracts it from the original image with minimal erosion of the target feature, which is critical in the estimation of fluorescence signals within the cell boundary (23).

### 2.6. Quantitative PCR

Quantitative PCR (qPCR) was performed using the MIC qPCR system (BioMolecular Systems, Upper Coomera, QLD, Australia). Each reaction was prepared using 2× BX-Master Mix (Cat. # BNROP-0049, NICSRO™) according to the manufacturer’s instructions. Primer sequences targeting the *vanA, vanB*, and the V3-V4 regions of 16S rRNA are listed in Table 1. Using the V3-V4 region as a reference, the relative quantification cycle (ΔC_q_) was calculated as the difference between the C_q_ of the target gene and the C_q_ of the reference, and ΔΔC_q_ was calculated as the difference in ΔC_q_ between conditions.

**Table 1.**
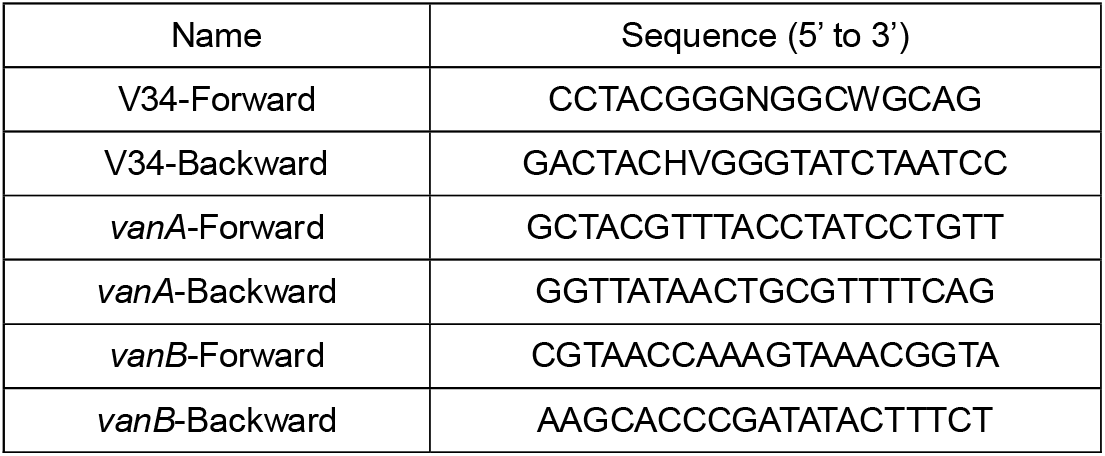
Sequences of qPCR primers used for the amplification of *vanA, vanB*, and the V3-V4 regions of 16S rRNA.

## 3. RESULTS

### 3.1. Detection and differentiation of vanA and vanB genes

To detect low-abundance mRNAs, PNA probes with high affinity and specificity to the target sequence are required. Previously, we distinguished bacterial species by using PNA probes targeting the variable regions of 16S rRNA that are unique to each species but conserved across different strains of the same species (24). Following the same design principle, conserved regions among D-Ala-D-Lac ligases, including *vanA* and *vanB*, as well as *vanB*-specific regions, were identified using the Comprehensive Antibiotic Resistance Database (CARD) (Fig. 1A, Materials and Methods). 15-bp-long candidate sequences were screened based on purine content, GC content, purine stretch length, self-complementarity, and off-target binding potential to other genes or 16S rRNA. A universal probe targeting both *vanA* and v*anB* (van-Univ) was designed to detect resistance due to either operon (Table 1). To identify the operon responsible for the resistance, a set of probes (vanB-1, vanB-2, vanB-3) targeting unique sequences of *vanB* was designed. These probes were designed to have at least two mismatches to any other regions in the entire *E. faecium* genome (Table 2). A previously validated PNA probe targeting a conserved region in the 16S rRNA of *E. faecium* (PNA-16S) was used to identify *E. faecium* cells. *vanA*-genotype (NCCP S16; VRA), *vanB*-genotype (NCCP 11522; VRB), and vancomycin-susceptible (KCTC 13225; VSE) *E. faecium* strains were used for tests. The antibiotic resistance of VRA and VRB was confirmed by qPCR and planktonic growth assay (Fig. 1B, Fig. S1).

**Table 1.**
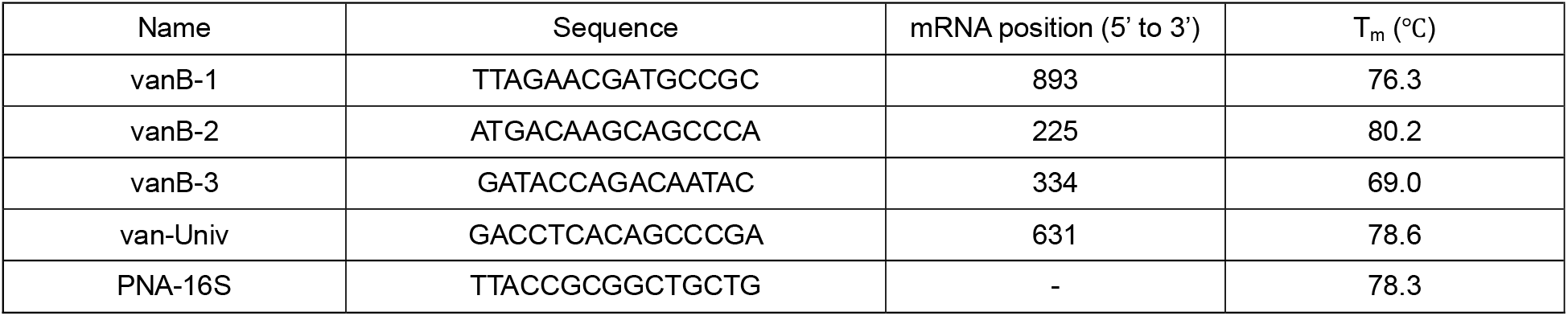
The sequences of the PNA-FISH probes. Positions were assigned based on the *vanA* and *vanB* mRNA sequences from CARD. The melting temperatures (T_m_) were calculated based on PNA-DNA hybrids; the corresponding PNA–RNA hybrids are expected to have slightly higher T_m_ values.

**Table 2.**
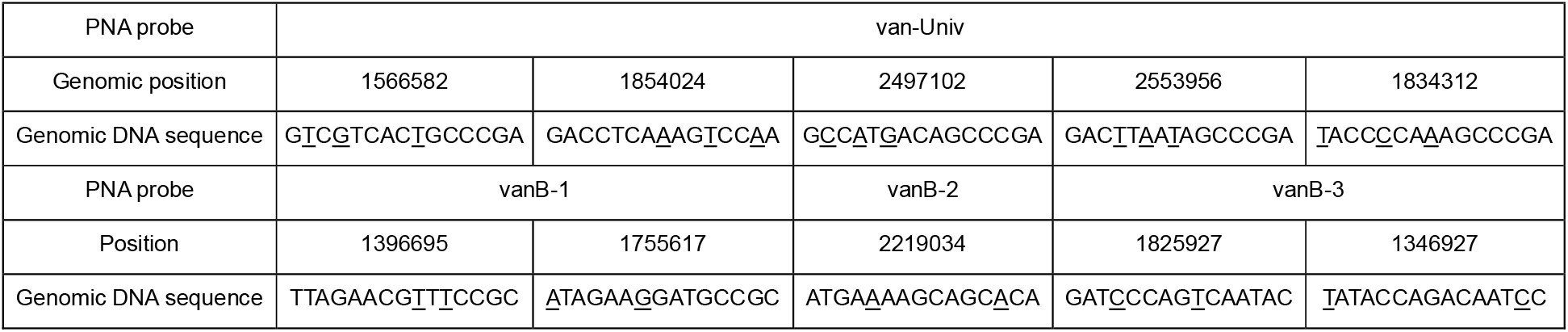
Off-target sites of the PNA probes. Potential off-target sites on *E. faecium* genome, GCF_009734005.1_ASM973400v2, with the least number of mismatches are shown. Mismatch positions are underlined.

**Figure 1.**
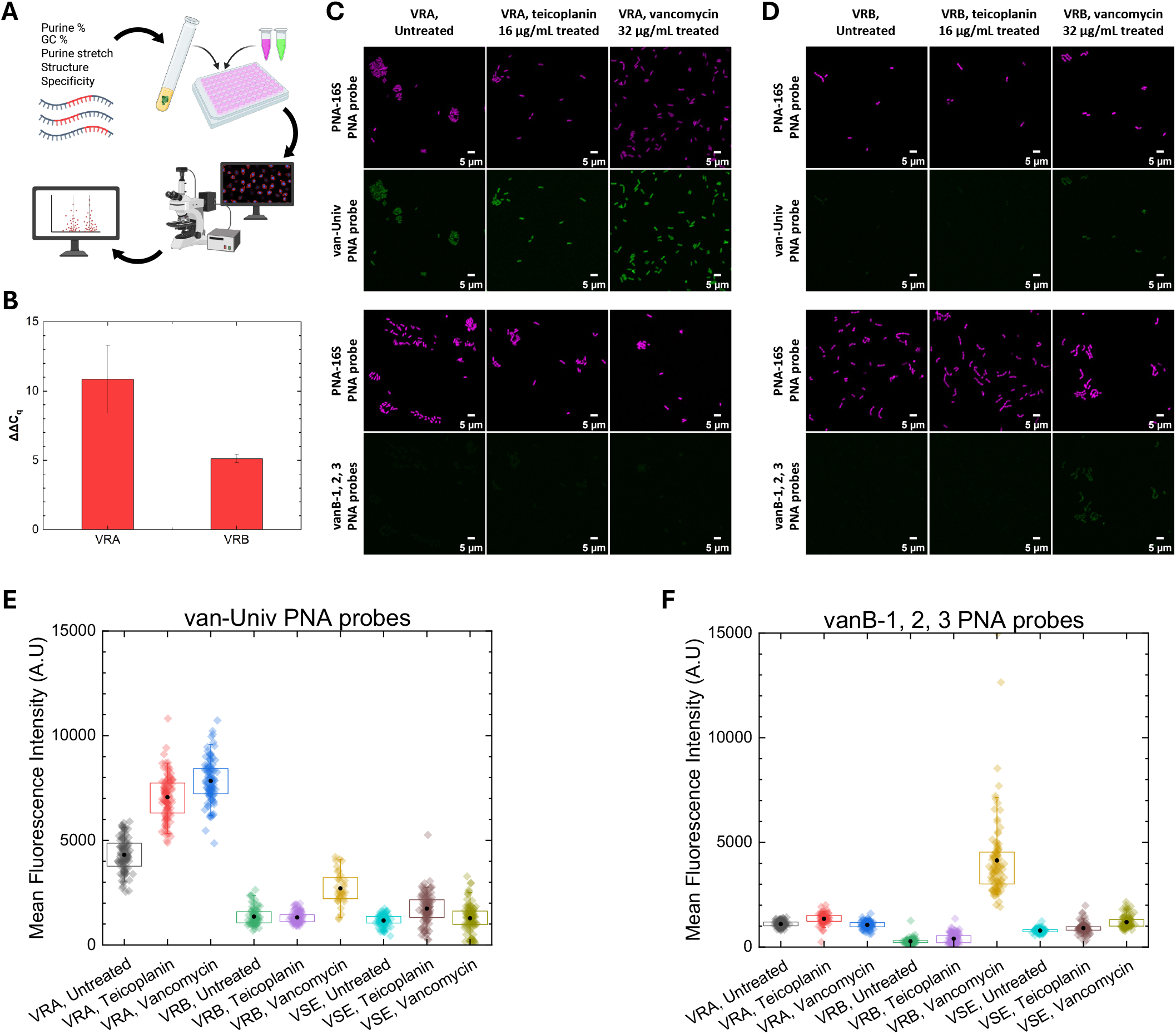
Detection and discrimination of vancomycin-resistant *E. faecium* using PNA-FISH targeting vancomycin-resistance-associated mRNA. **(A)** Schematic procedure to design PNA probes, stain bacterial cells, acquire fluorescence images, and analyze data. Target regions for PNA probe binding were selected based on five criteria; purine content, GC content, length of consecutive purine stretches, self-complementarity, and sequence specificity. **(B)** qPCR analysis showing differential expression of *vanA* and *vanB* mRNA in VRA and VRB, respectively, following exposure to 32 μg/mL of vancomycin. Mean ΔC_t_ values and errors from triplicate measurements are shown. **(C, D)** Representative PNA-FISH images of VRA (C) and VRB (D) strains without antibiotic treatment, with 16 μg/mL teicoplanin, or 32 μg/mL vancomycin for 1 hr. Scale bar, 5 μm. **(E, F)** Mean fluorescence intensities of individual VRA, VRB, and VSE cells with or without antibiotic treatment. Each colored dot represents the mean fluorescence intensity within a single cell. Black dots indicate mean values, and boxes represent the interquartile range (25th–75th percentiles).

Bacterial cells from each strain were treated with antibiotics prior to immobilization on a PLL-coated plate (Materials and Methods). After fixation and permeabilization, cells were hybridized with fluorescently labeled PNA probes. To quantify ARG-specific FISH signals from individual bacterial cells, two-color confocal imaging was performed. van-Univ and vanB-1, 2, 3 were labeled with Cy3, and PNA-16S was labeled with Cy5. Fluorescence images from PNA-16S were used to define cell boundaries and generate segmentation masks, which were then applied to the Cy3 channel to quantify the total fluorescence intensity of van-Univ or vanB-1, 2, 3 in individual cells.

VRA cells exhibited strong baseline fluorescence signals from van-Univ, which further increased upon treatment with teicoplanin or vancomycin (Fig. 1C). In contrast, VRB cells exhibited relatively lower fluorescence signals, comparable to those of VSE cells, which increased upon vancomycin treatment, but not upon teicoplanin treatment, consistent with the known lack of *vanB* induction by teicoplanin (Fig. 1D). To distinguish between VRA and VRB strains, cells were hybridized with vanB-1, 2, 3 and imaged (Fig. 1C, D). VRA cells exhibited no significant difference in fluorescence compared to VSE cells and no increase upon antibiotic treatment (Fig. 1C). In contrast, VRB cells exhibited a substantial increase in fluorescence exclusively upon vancomycin treatment (Fig. 1D). Mean fluorescence intensities of individual cells were analyzed to examine the distribution of signals among cells (Materials and Methods, Fig. 1E, F). The discrimination accuracy was calculated by finding a fluorescence threshold that equalizes the percentage of target cells (VRA or VRB) above it and the percentage of non-target cells (VSE) below it. van-Univ probe discriminated VRA from VSE with 97.94% accuracy even without antibiotic treatment, and it further increased to 99.96% and 99.99% with teicoplanin and vancomycin treatment, respectively (Fig. 1E). vanB-1, 2, 3 probes discriminated vancomycin-treated VRB cells from vancomycin-treated VSE cells with 95.64% accuracy (Fig. 1F). These results demonstrate distinct transcriptional responses of *vanA* and *vanB* operons to teicoplanin and vancomycin, as well as the capability of PNA probes to detect antibiotic resistance at the single-cell level.

### 3.2. Evaluation of individual vanB PNA probes

While vancomycin resistance was effectively detected using the universal probe (van-Univ), which targets conserved regions across the *vanA* and *vanB* operons, a set of three probes was employed for specific detection of *vanB* gene in the above experiments. To evaluate the hybridization efficiency of each probe, the vanB probes were tested individually. All probes produced detectable signals in VRB cells (Fig. 2A, B); however, the vanB-3 probe exhibited weaker fluorescence compared to the others. The mean fluorescence intensity of vanB-3 was approximately half of that of vanB-1 or vanB-2 (Fig. 3B). The combined signals obtained using all vanB probes were expected to approximate the sum of the signals from individual probes. However, individual probes exhibited lower fluorescence intensity when used alone than in combination. This observation suggests that the simultaneous application of multiple probes may induce cooperative binding, potentially by altering the secondary structure of the target mRNA, thereby increasing target accessibility.

**Figure 2.**
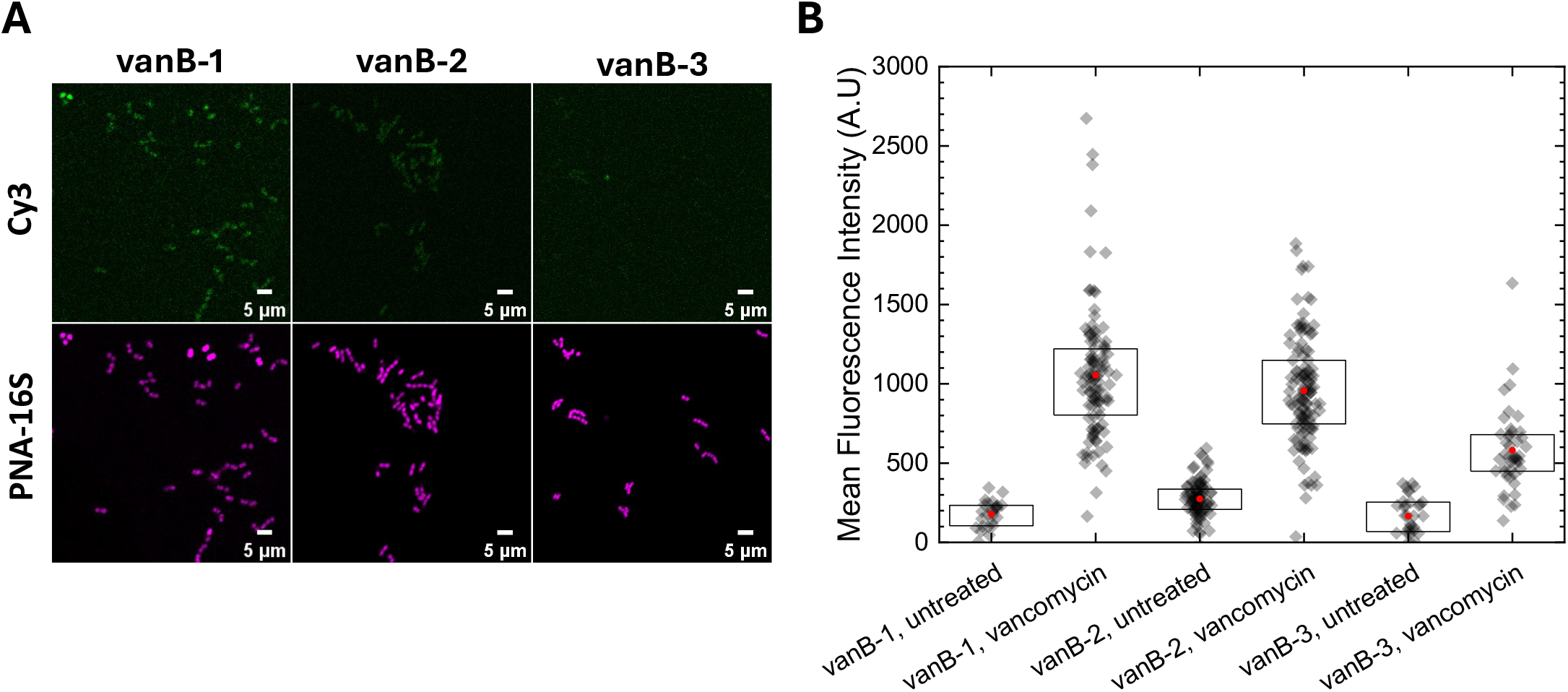
Evaluation of individual vanB PNA probes. **(A)** Representative PNA-FISH images of VRB cells hybridized with vanB-1, vanB-2, or vanB-3. Fluorescence images from the PNA-16S channel were used for cell identification. The contrast of the Cy3 channel was increased twofold relative to Figures 1 and 2 for visualization purposes. **(B)** Mean single-cell fluorescence intensities for each probe, with or without vancomycin treatment. Boxes indicate the interquartile range (25th–75th percentiles).

**Figure 3.**
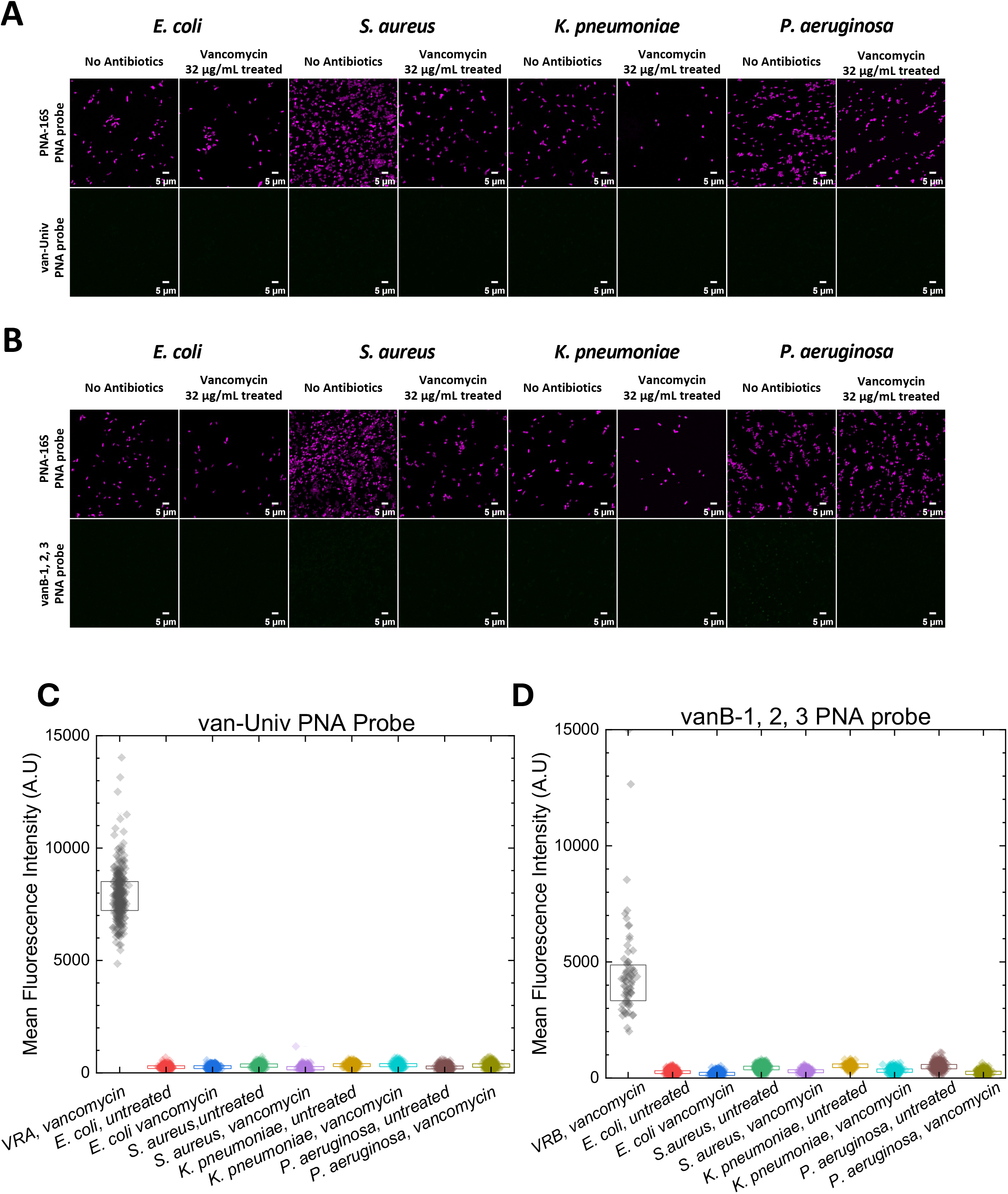
Validation of probe specificity for non-target bacterial species. **(A, B)** Representative PNA-FISH images of each non-target species hybridized with van-Univ (A) or vanB-1, 2, 3 (B), together with PNA-16S for cell identification. **(C, D)** Mean single-cell fluorescence intensities for each species, with or without vancomycin treatment. Boxes indicate the interquartile range (25th–75th percentiles).

### 3.3. Specificity of PNA probes for VREfm

To extend this technique to clinical applications, where samples typically contain a mixture of microbial species, it is essential to evaluate potential nonspecific binding of the probes. Four major bacterial species associated with sepsis—*Escherichia coli, Staphylococcus aureus, Klebsiella pneumoniae*, and *Pseudomonas aeruginosa—*were tested for nonspecific binding of van-Univ and vanB-1, 2, 3. None of the tested strains contained vancomycin resistance genes, and the absence of *vanA* and *vanB* was confirmed by whole-genome sequencing and PCR.

Individual bacterial cells were successfully detected using the PNA-16S probe, while neither van-Univ nor vanB-1, 2, 3 produced detectable fluorescence signals in these cells (Fig. 3A, B). The fluorescence intensity distributions of individual cells were clearly separated from those of vancomycin-treated VRA or VRB across all tested species, regardless of vancomycin treatment. The mean fluorescence intensity of these non-target species was at least nine-fold lower than that of vancomycin-treated VRA or VRB (Fig. 3C, D). These results demonstrate the high specificity and robustness of the PNA-FISH probes in distinguishing *vanA*- and *vanB*-genotype *E. faecium* from other clinically relevant pathogens, supporting their potential as a rapid, culture-independent alternative to conventional molecular diagnostics.

### 3.4. Detecting antibiotic resistance by wide-field imaging

Although PNA probes targeting *vanA* and *vanB* genes produced strong fluorescence signals (Fig. 1C and D), these measurements were obtained using confocal microscopy, which is not usually available in clinical laboratories. To evaluate the feasibility of a more accessible imaging modality, fluorescence signals were analyzed using a wide-field fluorescence microscope. Raw wide-field images exhibited substantial background fluorescence, resulting in a low signal-to-background ratio (SBR) of 1.1–1.2. Application of a background elimination algorithm substantially improved image contrast, increasing the SBR to 7.2 (Fig. 4A). Quantitative analysis of mean fluorescence intensity at the single-cell level enabled discrimination of vancomycin-treated VRB from vancomycin-treated VSE, achieving a discrimination accuracy of 93.4% (Fig. 4B).

**Figure 4.**
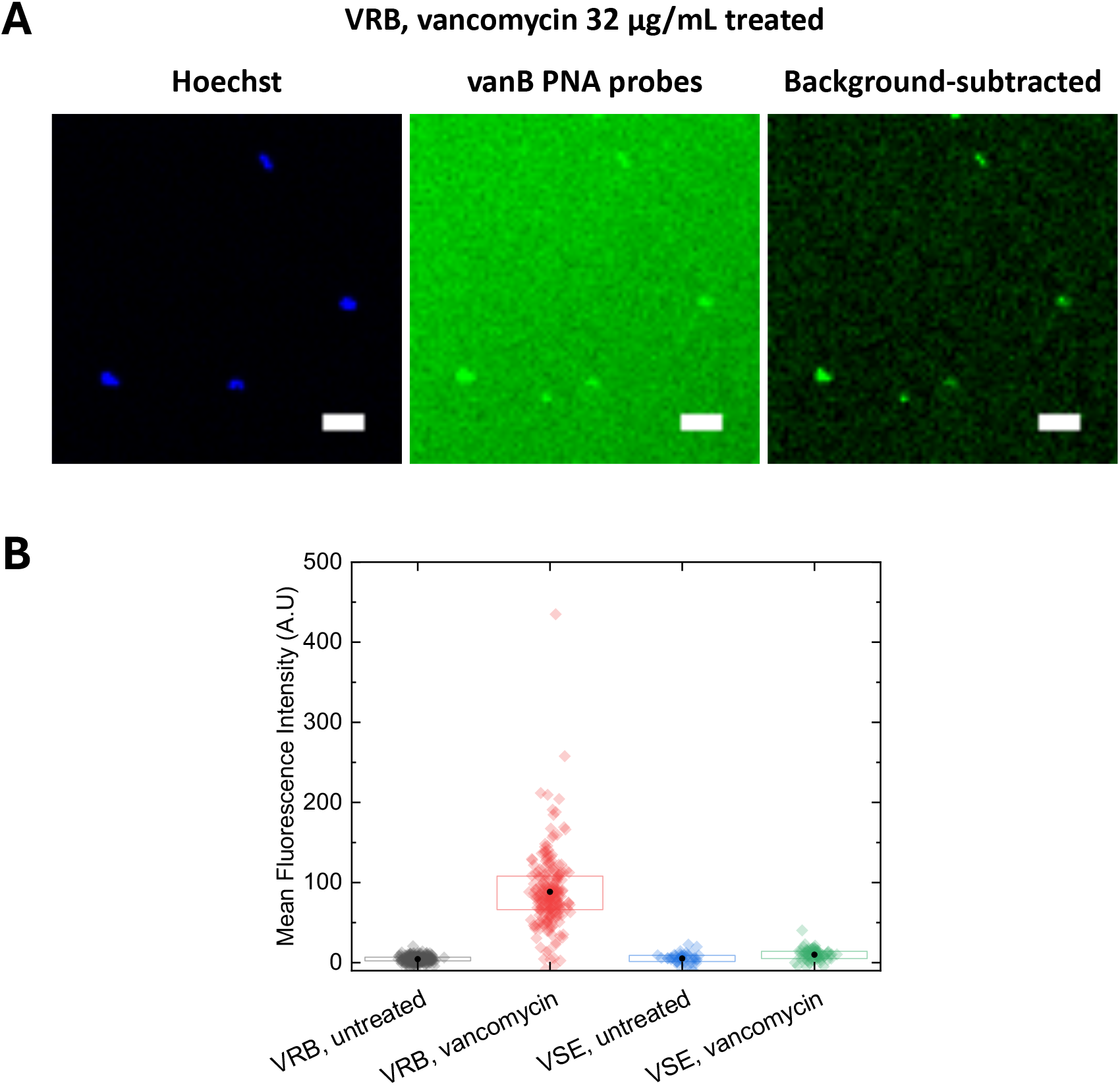
Wide-field fluorescence imaging of PNA-FISH. **(A)** Representative wide-field fluorescence images of VRB cells stained with Hoechst and hybridized with vanB-1, 2, 3 probes. Scale bars, 10 µm. **(B)** Mean single-cell fluorescence intensities measured from background-subtracted Cy3 channel images. Black dots indicate mean values for each condition, and boxes represent the interquartile range (25th–75th percentiles).

## 4. CONCLUSIONS

This study demonstrates the potential of PNA-FISH as a rapid, culture-independent diagnostic approach for detecting vancomycin resistance in VREfm. We developed a strategy for selecting optimal target regions within mRNA to improve probe efficiency. The distinct fluorescence responses of *vanA* and *vanB* genes to teicoplanin and vancomycin treatment enabled reliable differentiation of resistance genotypes at the single-cell level. The absence of detectable signals in non-target bacterial species commonly found in bloodstream infections further supports the specificity of the probes for clinical applications. In addition, the feasibility of detection using wide-field fluorescence microscopy highlights the potential for broader implementation in standard laboratory settings. Multiplex detection using spectrally distinct PNA probes further enables simultaneous identification of multiple antibiotic resistance genes. Together, these results establish PNA-FISH as a rapid, amplification-free, and scalable platform for antimicrobial resistance diagnostics.

## DATA AVAILABILITY

Data will be made available upon request.

